# Beneficial *Lactiplantibacillus plantarum* promote Drosophila growth by down-regulating the expression of PGRP-SC1

**DOI:** 10.1101/2021.07.16.452638

**Authors:** Marialaura Gallo, Justin M. Vento, Pauline Joncour, Andrea Quagliariello, Elisa Maritan, Chase L. Beisel, Maria Elena Martino

## Abstract

Animals and their commensal bacteria are known to reciprocally influence many traits of their physiology. Specifically, microbes contribute to the maintenance of the immune system homeostasis, improve host digestive processes, and sustain host growth and development. Several studies have reported that such effects result from an intricate network of nutritional, metabolic and immune inputs and partly rely on the capacity of microbes to regulate the host’s transcriptional response. However, these evidences mainly come from comparing the transcriptional response caused by commensal bacteria with that of axenic animals, making it difficult to identify the specific animal genes that are regulated by beneficial microbes. Here, we employ a well-established model of nutritional symbiosis, *Drosophila melanogaster* associated with *Lactiplantibacillus plantarum*, to understand the host genetic pathways regulated by beneficial bacteria and leading to improved host growth and development. Using isogenic *L. plantarum* strains bearing different growth-promoting effects, we show that the microbial benefit to the host relies on the down-regulation of peptidoglycan- recognition proteins. In particular, we report that the lower expression of PGRP-SC1 exerted by growth-promoting bacteria is responsible for their higher proliferation and the consequent increased production of beneficial metabolites, which ultimately leads to improved host growth and development. Our study helps elucidate the mechanisms underlying the beneficial effect exerted by commensal bacteria, defining the role of PGRP-SC1 in the relationship between Drosophila and its gut microbes.

**IMPORTANCE:** Commensal bacteria are in constant association with their animal hosts, significantly affecting animal physiology through an intricate network of nutritional, metabolic and immune inputs. Yet, how beneficial bacteria specifically improve animal health is not fully understood. Here, we used a well-established model of nutritional symbiosis to understand how beneficial gut microbes improve host growth via regulation of its transcriptional response. Our study advances the current knowledge in host-microbe interactions by demonstrating that commensal bacteria improve fly growth by actively regulating the expression of immune effectors, which lead to higher immune tolerance. This leads to higher bacterial proliferation and the increased production of beneficial microbial metabolites, which are then consumed by the host. Our results shed light on the complex mechanisms underlying the relationships between a host and its gut microbes.

## INTRODUCTION

Metazoans live in constant association with microbial communities, and their reciprocal interaction is known to favour many traits of both partners’ physiology (1, 2). On one hand, animal hosts can benefit the growth and metabolism of their microbial partners, specifically through the secretion of a complex blend of metabolites (3–5). On the other hand, beneficial bacteria sustain animal health by contributing to the maintenance of the immune system homeostasis (6), confer protection against pathogen infections (7, 8), promote intestinal epithelium renewal (9, 10), and influence host lifespan and development (11–13). In addition, the gut microbiota bolsters the host digestive processes through the production of a large collection of enzymes that favour the metabolism of dietary molecules as well as specific organic compounds such as vitamins, amino acids and short-chain fatty acids (5, 14). This type of relationship is referred to as nutritional symbiosis (15). Most insects undertake facultative nutritional symbioses with their microbial communities (16, 17), where both partners benefit from each other’s presence without relying on one another for their survival (18). *Drosophila melanogaster*, associated with its microbiota, is a well-known example of such a relationship (19, 20) and represents a powerful model to study host-microbe interactions (21, 22). Wild and lab-reared Drosophila are associated with relatively simple beneficial bacterial communities. These communities mainly include *Acetobacteraceae* and *Lactobacillaceae* families, with *Acetobacter pomorum and Lactiplantibacillus plantarum* as the most common species of the *Drosophila melanogaster* microbiota (20, 23, 24). *Drosophila* microbiota influences the host’s physiology throughout its entire life (19, 20, 22). Beside the above mentioned benefits, fly gut microbes are also known to impact their host’s physiology via regulation of many host transcriptional pathways (25). Numerous studies have been conducted to characterize the impact of the microbiota on the gene expression of the adult fly. Commensal bacteria are involved in the modulation of intestinal immune homeostasis (26–29), they shape gut morphology through enhancement of stem cell proliferation and epithelial renewal rate (10, 25, 30), and they have also been shown to mediate transgenerational inheritance of responses to environmental changes (31, 32). In addition, it has been demonstrated that gut microbes influence host phenotype by promoting the coexpression of groups of genes involved in major signalling pathways, among which are the response of larval growth, adult physiology and nutrient availability (33).

One of the most striking beneficial effects exerted by gut microbes is host growth promotion (34–36). Some bacterial species associated with *D. melanogaster* have been demonstrated to be able to promote fly growth and maturation rate during the juvenile phase in poor-nutrient conditions (34). The growth-promoting ability of the gut microbiota is strain-dependent and is exerted through the enhancement of the systemic production of two hormonal growth signals: Ecdysone (Ecd) and Drosophila insulin-like peptide (dILP) via increased TOR kinase activity in both fat body and prothoracic gland (34, 37). In addition, it has been shown that fly growth promotion mediated by gut microbes results, at least in part, from the upregulation of digestive enzyme expression in the midgut via IMD/Relish activity (37). These insights came from comparing Drosophila with a microbiota (either single-strain associations or complex communities) versus without a microbiota (germ-free condition) (25, 28, 29, 33, 37, 38), and they largely contributed to our knowledge of the molecular mechanisms underlying the role of gut microbes in host physiology. Nevertheless, not all bacterial species or strains associated with Drosophila microbiota are beneficial. As a consequence, a clear understanding of the host genetic pathways that are specifically regulated by beneficial bacteria remains elusive.

We recently demonstrated that a strain of *L. plantarum* (*Lp*^NIZO2877^) bearing a moderate *Drosophila* growth-promoting ability was able to improve its beneficial effect by adapting to the fly nutritional environment (39). The *Lp*^NIZO2877^-evolved strain (*Lp*^Fly.G2.1.8^) showed a single mutation in the acetate kinase A (*ackA*) gene, causing the increased production of N-acetylated amino acids, which were sufficient to improve *L. plantarum* growth-promoting capacity. In this study, we coupled comparative transcriptome analyses and CRISPR-Cas-based microbial engineering on such isogenic strains to reveal the specific host transcriptional pathways regulated by the association with beneficial gut bacteria and leading to enhanced animal growth. The presence of a single genetic difference between the bacterial strains, which is responsible for the improvement in growth promotion, allowed us to uniquely ascribe the differential host transcriptional response to the microbial benefit. We observe that growth-promoting *L. plantarum* strains are able to specifically lower the expression of the *Drosophila* peptidoglycan recognition protein (PGRP) SC1, whose activity is regulated by *L. plantarum ackA* gene function and its metabolic products. In addition, we demonstrate that the down-regulation of *PGRP-SC1* gene in larvae associated with a mid-beneficial strain is sufficient to recapitulate the beneficial effect of the strong growth promoting strain and that such benefit is directly linked to bacterial growth.

## RESULTS

### Commensal bacteria significantly affect *Drosophila* transcriptional response

To unravel the *Drosophila* transcriptional pathways specifically regulated by beneficial gut microbes, we compared the transcriptome of *D. melanogaster* larvae during the second instar associated with two strains of *L. plantarum* promoting Drosophila growth to different extents (*Lp*^NIZO2877^ *and Lp*^FlyG2.1.8^, gnotobiotic larvae) with germ-free (GF) larvae as negative control. *Lp*^NIZO2877^ is a *L. plantarum* strain isolated from processed human food (40) showing a moderate growth promotion both in *Drosophila* and in mice (35). *Lp*^FlyG2.1.8^ is a *Lp*^NIZO2877^-derived strain experimentally evolved in poor-nutrient conditions in the presence of *D. melanogaster*. *Lp*^FlyG2.1.8^ is isogenic to *Lp*^NIZO2877^, with one exception: *Lp*^FlyG2.1.8^ bears a tri-nucleotide deletion in the *ackA* gene, which is responsible for a significant improvement in larval growth promotion compared to its ancestor (39). The rationale behind the choice of such strains is that any differences in host transcript levels could be directly ascribed to the single bacterial genetic mutation and, thus, to the improved growth promoting effect. Following bacterial association with GF embyos, we performed Drosophila RNA extraction on size-matched larvae (2.4 mm length). Specifically, considering that microbiota-associated larvae develop faster than axenic larvae (34), we collected the *Lp*-associated larvae 4 days after monoassociation, while the GF larvae were collected after 5 days (Fig. S1). The transcriptome sequencing of Drosophila larvae yielded 47-31 million reads for all replicate samples. We analysed the transcripts using a dedicated R script and obtained a data set of 17,560 genes (Table S1). Among these genes, 559 genes were differentially expressed between axenic and gnotobiotic larvae that satisfied our criteria for significant differential expression (p-value < 0.05 and −1.5 to +1.5 fold). 285 genes were differentially expressed in both gnotobiotic conditions (*Lp*^NIZO2877^- and *Lp*^FlyG2.1.8^-associated larvae) compared to GF larvae, among which 106 resulted to be up-regulated and 179 down-regulated (Table S1). Next, we analyzed the remaining 274 genes and we observed that 209 were differentially expressed between *Lp*^FlyG2.1.8^-associated larvae and GF larvae, with 96 up-regulated and 113 down-regulated genes, whereas the other 65 genes were differentially expressed between *Lp*^NIZO2877^-associated larvae and GF larvae, with 22 up-regulated and 43 down-regulated (Fig. S2A, Table S1).

We assigned biological pathways to the identified 559 genes using DAVID Gene Functional Classification Tool (41), which annotates each gene and identifies the most relevant biological terms associated with a given gene. To improve the usefulness of the functional annotation analysis, we carried out DAVID clustering analysis for a group of up- and down-regulated genes, respectively, and we considered only the genes that met our statistical criteria (*Benjamini-adjusted-p-value* < 0.1).

The transcripts enriched in gnotobiotic larvae compared to GF larvae were dominated by the expression of lysozyme genes (i.e., *LysB, C, D* and *E*) involved in the hydrolysis of beta-linkages between N-acetylmuramic acid and N-acetyl-D-glucosamine residues in peptidoglycan, and the expression of heat shock protein (*Hsp*) genes (i.e., *Hsp22*, *23, 26, 27* and *28*) commonly expressed in response to stress resistance and adaptation (Fig. S2B, Table S2). Furthermore, gnotobiotic larvae showed significantly higher expression of immunity-related genes, including genes involved in peptidoglycan (PGM) recognition and the negative regulation of innate immune response, such as *PGRP-SB1*, *PGRP-SD*, *PGRP-LA*, *PGRP-SCs* (Tables S1, S2). The genes down-regulated in gnotobiotic larvae compared to germ-free conditions were implicated in membrane function through the involvement of genes with catalytic activity, such as *Pkn* and *Adgf-A2* (Tables S1, S2).

By comparing the transcriptional response between single-strain associated larvae and GF larvae, the GO term analysis revealed an enrichment of expression in genes implicated in DNA methylation (*Mt2*), DNA binding, and DNA-directed RNA polymerase activity (*Rpb5, RpI12*) in *Lp*^FlyG2.1.8^-associated larvae. In addition, we detected a significant decrease in the expression of genes involved in cell biogenesis, such as *l(2)gl*, the *crb* gene essential for the development of polarized epithelia and centrally involved in cell polarity establishment, and the *scrib* gene required for polarization of the embryonic imaginal disk and follicular epithelia (Fig. S2C, Table S2).

### Beneficial *L. plantarum* strains trigger downregulation of the peptidoglycan-recognition protein SC1 compared to mid-beneficial strains

Next, we sought to identify the host genes regulated by a beneficial bacterium, specifically improving fly growth. To do this, we compared the transcriptome profiles between *Lp*^NIZO2877^- and *Lp*^FlyG2.1.8^-associated larvae. Our analysis identified 21 genes differential expressed between the conditions (p < 0.05 and −1.5 to +1.5 fold), with 6 and 15 genes being down- and up-regulated in larvae associated with *Lp*^FlyG2.1.8^-associated larvae compared with *Lp*^NIZO2877^-associated larvae, respectively (Fig. 1, Table S3). The transcripts found to be enriched in larvae associated with the beneficial strain *Lp*^FlyG2.1.8^ were dominated by the expression of the Sperm-Leucylaminopeptidase 3 (*S-Lap3*) gene (22.8 fold), a major constituent of paracrystalline material from *D. melanogaster* sperm involved in the fertility of male flies (42). We also observed the up-regulation of genes involved in the production of larval cuticle protein (*Lcp4* and *Lcp65Ab1),* proteolysis (*CG43124* and *CG43125*), signal peptides (*CG14258, CG14259* and *CG15597*), the odorant-binding protein 56e (*Obp56e*) involved in nutrient sensing and methuselah-like 6 (*mthl6)* implicated in Drosophila development and lifespan. At the same time, *Lp*^FlyG2.1.8^-associated larvae showed down-regulation of genes involved in chitin metabolism (*thw*), the Niemann-Pick type C-2c (*Npc2c*) gene, implicated in sterol transport, and two signal peptides (*CG11381, CG7465*) (Table S4). Surprisingly, we found that *Lp*^FlyG2.1.8^*-*associated larvae showed a lower expression of *PGRP-SC1a* and *PGRP-SC1b* genes compared to *Lp*^NIZO2877^-associated larvae. No difference was detected in the regulation of other PGRPs (Table S1, S4). The specific lower expression of *PGRP-SC1* in larvae associated with the strong growth-promoting strain *Lp*^FlyG2.1.8^ prompted us to investigate the causes of such immune system down-regulation.

**FIG 1.**
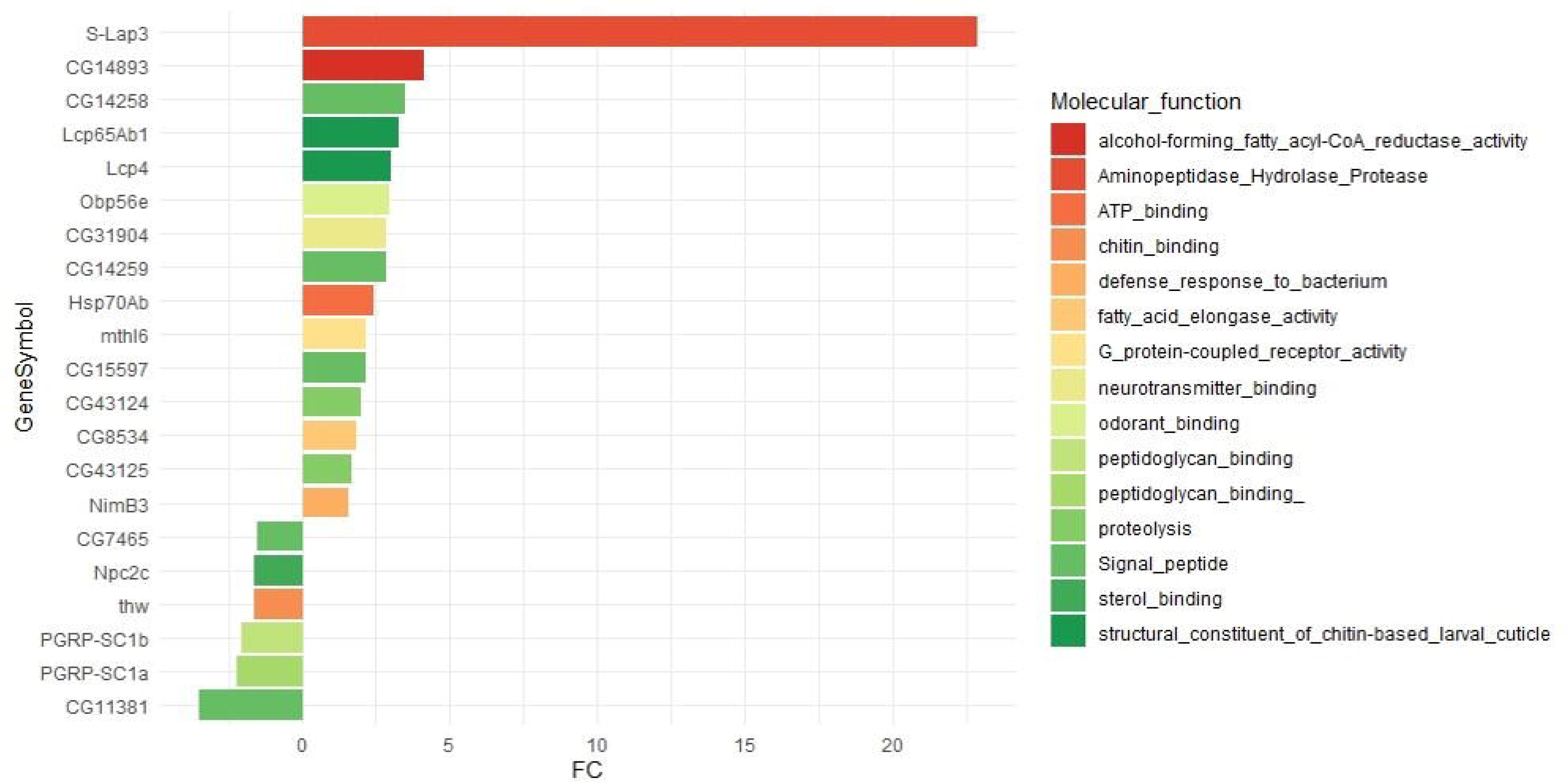
*Lp*^FlyG2.1.8^ significantly alter Drosophila transcriptional response compared to *Lp*^NIZO2877^. Bar chart of the genes with significantly differential expression (up- and down-regulated) between *Lp*^FlyG2.1.8^- and *Lp*^NIZO2877^-associated larvae (p < 0.05 and −1.5 to +1.5 fold). Each colour refers to the molecular function associated with each gene.

### *Drosophila PGRP* expression relies on *L. plantarum ackA* function

First, to validate the RNA-Seq data, transcript abundance of *PGRP-SC1* genes was assessed on size-matched larvae newly associated with *Lp*^NIZO2877^ and *Lp*^FlyG2.1.8^ strains. The amplification results confirmed significantly lower expression of *PGRP-SC1* in *Lp*^FlyG2.1.8^-associated larvae (Fig. 2A). As *Lp*^FlyG2.1.8^ has been experimentally co-evolved with the fruitfly, we hypothesized that the down-regulation of *PGRP-SC1* expression might be due to mechanisms of immune tolerance. To test this hypothesis, we analysed the expression of *PGRP-SC1* in larvae associated with *Lp*^DietG20.2.2^, a different *Lp*^NIZO2877^-derived strain experimentally evolved on the fly diet and bearing a mutation in the same *ackA* gene as *Lp*^FlyG2.1.8^ that also improved growth promotion (39). *Lp*^DietG20.2.2^ led to the down-regulation of *PGRP-SC1* compared to *Lp*^NIZO2877^-associated larvae (Fig. 2A), thus suggesting that *PGRP-SC1* expression was independent of the strain evolutionary history. Next, we focused on the microbial molecular mechanisms that led to the down-regulation of *PGRP-SC1* expression. Specifically, both *Lp*^FlyG2.1.8^ and *Lp*^DietG20.2.2^ strains bear non-synonymous mutations in the *ackA* gene which have been predicted to result in the loss of function of the gene (39). We thus asked whether the bacterial *ackA* was directly involved in the regulation of *PGRP-SC1*. To investigate this, we employed CRISPR/Cas9-based bacterial genetic engineering (43) to knock out the *ackA* gene in the ancestor strain *Lp*^NIZO2877^. *Lp*^*ΔackA*^ showed a significantly improved growth-promoting effect compared to the ancestor *Lp*^NIZO2877^ (p < .0001) (Fig. 2B). No significant difference was found when comparing *Lp*^*ΔackA*^ and *Lp*^FlyG2.1.8^ and *Lp*^DietG20.2.2^ strains. These results demonstrate that the loss-of-function of the *L. plantarum ackA* gene is sufficient to improve growth promotion by *L. plantarum* in Drosophila.

**FIG 2.**
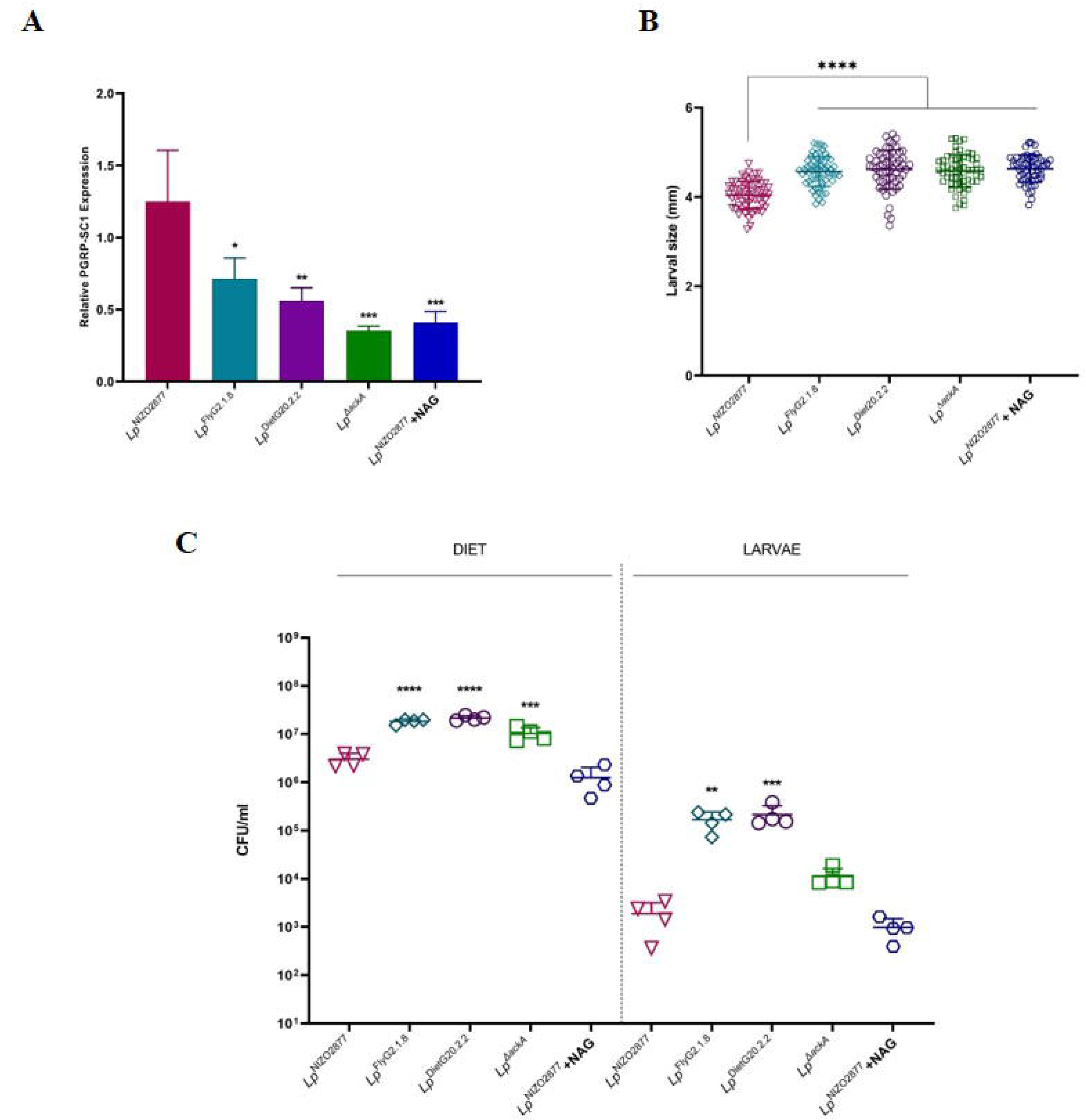
PGRP-SC1 down-regulation by beneficial *L. plantarum* strains leads to higher bacterial colony counts and improved fly growth promotion. (A) Relative expression of PGRP-SC1 gene obtained by performing a qRT-PCR analysis on the transcriptome of *Drosophila* larvae mono-associated with the bacterial strain *Lp*^NIZO2877^, *Lp*^FlyG2.1.8^, *Lp*^DietG20.2.2^, *Lp*^ΔackA^ mutant, and *Lp*^NIZO2877^ supplemented with N-acetyl glutamine (NAG), respectively. Lines above each bar represent the mean with the standard deviation (SD) calculated by analyzing five biological replicates per condition. Normalization was performed using the housekeeping gene *rp49*. All conditions were compared to *Lp*^NIZO2877^ by performing One-way ANOVA test (*p≤0.05, **p≤0.01, ***p≤0.001). (B) Longitudinal size (mm) of *Drosophila* larvae measured after 7 days of incubation with the bacterial strain *Lp*^NIZO2877^, *Lp*^FlyG2.1.8^, *Lp*^DietG20.2.2^, *Lp*^ΔackA^, and *Lp*^NIZO2877^ supplemented with N-acetyl glutamine (NAG), respectively. Each symbol refers to the larval size obtained from one out the N≥60 *Drosophila* larvae analyzed for each condition, with bars referring to the respective mean and standard deviation (SD). All conditions were compared to *Lp*^NIZO2877^ by performing One-way ANOVA test (*p≤0.05, **p≤0.01, ***p≤0.001, ****p≤0.0001). (C) Microbial load (CFU/mL) of the bacterial strains Lp^NIZO2877^, Lp^FlyG2.1.8^, Lp^DietG20.2.2^, Lp^ΔackA^, and Lp^NIZO2877^ supplemented with N-acetylglutamine (NAG), after 4 days of incubation in the fly diet and in the intestine of Drosophila larvae, respectively. Each symbol represents one out of the four replicates analyzed for each condition, with bars indicating the respective mean and standard deviation (SD). All conditions were compared to Lp^NIZO2877^ by performing One-way ANOVA test (*p≤0.05, **p≤0.01, ***p≤0.001, ****p≤0.0001).

We then asked whether the *ackA* loss-of-function was responsible for the down-regulation of the immune response observed in *Lp*^FlyG2.1.8^-associated larvae. To test this, we analysed the expression of *PGRP-SC1* in larvae associated with *Lp*Δ*ackA*. Figure 2A shows that the absence of *ackA* resulted in down-regulation of the *PGRP-SC1* compared to *Lp*^NIZO2877^-associated larvae (Fig.2A). Next, we wondered whether the bacterial metabolic response resulting from the *ackA* mutation was directly involved in Drosophila immune regulation, or, instead, whether *PGRP-SC1* expression was governed by a more complex mechanism resulting from *ackA* mutation. Specifically, we previously demonstrated that the presence of the *Lp*^FlyG2.1.8^-*ackA* variant leads to an increase in the bacterial production of N-acetyl-glutamine (NAG), which is sufficient to recapitulate the improved beneficial effect exerted by *Lp*^FlyG2.1.8^ in the presence of its ancestor (39) (Fig. 2B). In light of this, we tested the effect of such a metabolite alone on Drosophila *PGRP-SC1* expression by supplementing NAG to Lp^NIZO2877^-associated larvae. Notably, we found that the presence of NAG was sufficient to down-regulate *PGRP-SC1* expression in Drosophila larvae (Fig. 2A). Taken together, our results demonstrate that *PGRP-SC1* expression in Drosophila is regulated by *L. plantarum ackA* function. Specifically, the *ackA* loss of function leads to an increase in the production of N-acetyl-glutamine, which is sufficient to cause *PGRP-SC1* down-regulation in Drosophila larvae.

### High expression of *PGRP-SC1* genes leads to lower microbial loads

Previous research has shown that *PGRP-SCs* act as negative regulators of IMD pathway activation. Their combined effect shapes the Drosophila antibacterial response and protects the fly from innocuous infections (27, 28, 44, 45). We thus wondered whether the higher expression of *PGRP-SC1* observed in *Lp*^NIZO2877^-associated larvae resulted in increased bacterial proliferation through negative regulation of the IMD pathway. To test this, we quantified the bacterial growth of *Lp*^NIZO2877^, *Lp*^FlyG2.1.8^, *Lp*^DietG20.2.2^, *Lp*^*ΔackA*^, and *Lp*^NIZO2877^ supplemented with NAG, both in the larval gut and in the fly diet at the same time point as the transcriptome analysis (4 days after association). In general, colony counts for *L. plantarum* were significantly lower in larval intestine compared to fly food (Fig. 2C). However, we observed that the colony counts for *Lp*^NIZO2877^ were always significantly lower compared to the other bacterial associations. The only exception was represented by the N-acetylglutamine addition, where *L. plantarum* load reached similar levels. Altogether, these data show that the expression of *PGRP-SC1* is inversely proportional to the growth of commensal bacteria.

### Down-regulation of *PGRP-SC1* is sufficient to improve Drosophila growth

So far, we have demonstrated that the loss of function of *L. plantarum ackA* gene is responsible for improving the bacterial growth promoting effect in Drosophila and that this happens through down-regulation of the fly *PGRP-SC1* gene. Following this result, we asked whether the down-regulation of *PGRP-SC1* was sufficient to improve larval growth. To test this, we used a mutant fly line bearing a deletion of the *PGRP-SC* gene (*PGRP-SC*^*Δ*^) (28). We mono-associated *PGRP-SC*^*Δ*^ GF embryos with *Lp*^NIZO2877^ and *Lp*^FlyG2.1.8^ strains and analysed their effect on larval growth compared to axenic conditions. Remarkably, we found that *Lp*^NIZO2877^ exerted the same beneficial effect as *Lp*^FlyG2.1.8^ in *PGRP-SC*^*Δ*^ larvae compared to GF conditions (Fig. 3A). At the same time, GF larvae were significantly smaller than bacterial-associated larvae. Following our previous results showing that *PGRP-SC1* expression is inversely proportional to bacterial growth (Fig. 2), we asked whether *Lp*^NIZO2877^ loads were higher in *PGRP-SC*^*Δ*^ flies than in wild type flies. Indeed, we observed that *Lp*^NIZO2877^ reached significantly higher colony counts, with comparable bacterial loads as those of *Lp*^FlyG2.1.8^ in both the mutant and wild type lines (Fig. 3B). Altogether, our results show that *L. plantarum* is able to improve Drosophila growth by regulating its immune response via the down-regulation of *PGRP-SC1*, which results in increased bacterial growth and proliferation.

**FIG 3.**
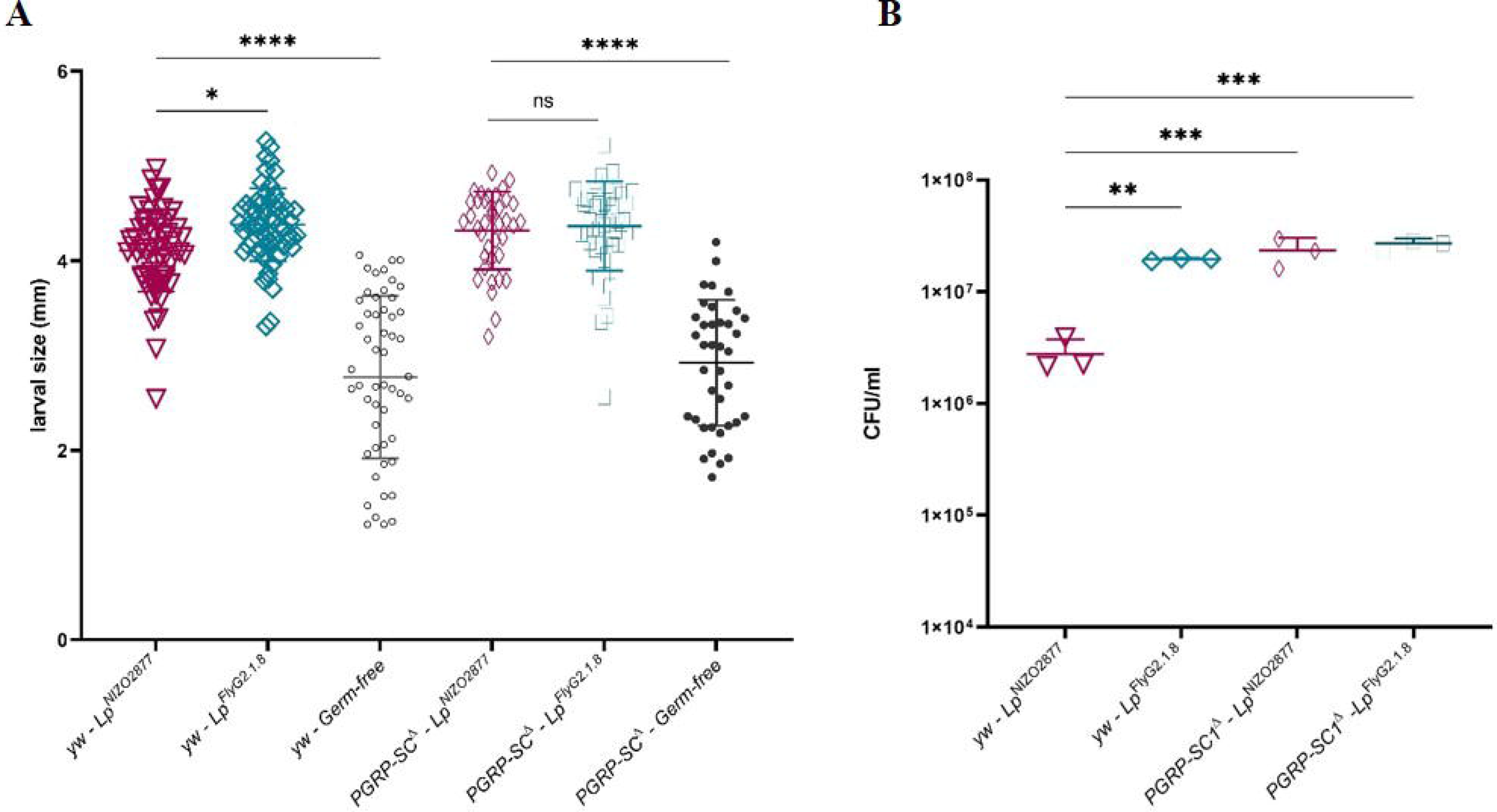
*Lp*^NIZO2877^ recapitulates the same growth-promoting effect of *Lp*^FlyG2.1.8^ in PGRP-SC1^Δ^ flies. Longitudinal size (mm) of Drosophila larvae performed on yellow-white (*yw*) wild-type population and on mutant fly larvae carrying a deletion of PGRP-SC1 gene cluster (PGRP-SC1^Δ^) both associated with the bacterial strains *Lp*^NIZO2877^, *Lp*^FlyG2.1.8^, respectively, or in germ-free (GF) condition, and measured after 7 days of growth. Each symbol refers to the larval size obtained from one out the N≥60 *Drosophila* larvae analyzed for each condition, with lines indicating the respective mean and standard deviation (SD). Each condition was compared with Lp^NIZO2877^-associated larvae of the respective fly background (*yw* or PGRP-SC1^Δ^) by performing One-way ANOVA test (*p≤0.05, **p≤0.01, ***p≤0.001, ****p≤0.0001). (B) Comparison among the microbial load (CFU/mL) of bacteria retrieved from the diet of yellow-white (yw) Drosophila larvae associated with the strains *Lp*^NIZO2877^, *Lp*^FlyG2.1.8^, *Lp*^Diet20.2.2^, *Lp*^ΔackA^, *Lp*^NIZO2877^ supplemented with N-acetyl glutamine (NAG), and PGRP-SC1^Δ^ Drosophila larvae associated with the strains *Lp*^NIZO2877^ and *Lp*^FlyG2.1.8^, respectively. Each symbol represents the mean value for each technical replicate, while lines are the mean obtained from the three values within the same condition with the respective standard deviation (SD). All conditions were compared to *Lp*^NIZO2877^ by performing One-way ANOVA test (*p≤0.05, **p≤0.01, ***p≤0.001, ****p≤0.0001).

## DISCUSSION

In the present study, we aimed at understanding the Drosophila transcriptional pathways regulated by beneficial gut commensals and leading to improved growth. Consistent with previous work, we showed that gut microbes strongly affect host’s gene expression (25, 33, 46), and such effects happen regardless of their benefit potential. Indeed, the presence of two isogenic *L. plantarum* strains differing in growth promotion significantly altered gene expression in Drosophila compared to axenic conditions. Analysis of the genes differentially expressed between gnotobiotic and axenic conditions revealed the up-regulation of genes related to host immune response in the presence of bacteria (Fig. S2, Tables S1, S2). It is well known that commensal bacteria are basal inducers of host immune responses (9, 22, 46–49). Our analysis highlighted the significant impact of peptidoglycan-recognition proteins (e.g., *PGRP-SB1*, *PGRP-SD*, *PGRP-LA*, *PGRP-SCs*) in Drosophila immune system regulation and response to bacteria. PGRPs represent the main and most diverse functional family of peptidoglycan detection proteins (50–55) and have been largely reported as crucial effectors in regulating the antibacterial response in Drosophila (56–58). We also detected an enriched expression of lysozyme genes (i.e., *LysB, C, D* and *E*), which are synthesized at a high rate in the midgut of larvae and adult flies and are known to have a specific role in the digestive process (59, 60). In addition, they are involved in the hydrolysis of beta-linkages between *N-acetylmuramic* acid and *N-acetyl-D-glucosamine* residues in bacterial peptidoglycans, likely supporting the action of the PGRP-SC genes in the antibacterial response (61). In addition to immune system regulation, bacterial association caused the up-regulation of genes known to be involved in stress response (e.g., *hsp23*, *GstD8*), proteolysis (*Jon66ci*, *Jon66cii*, *Jon44E*, *Jon99ci*, *Bace*, etc.) and metabolism (Fig. S2, Tables S1, S2). All these results are largely in agreement with those of previous studies examining the impact of gut commensals on *D. melanogaster* transcriptional regulation across different fly developmental stages and populations (46, 49).

We next sought to determine the host genes specifically altered by the presence of commensal bacteria that promote Drosophila growth and development. To do this, we compared the transcriptional response of Drosophila larvae associated with two isogenic *L. plantarum* strains: a strong growth promoting strain (*Lp*^FLYG2.1.8^), and the mid-beneficial strain *Lp*^NIZO2877^. We detected differential expression of only 21 genes between the two conditions. Specifically, *Lp*^FLYG2.1.8^ led to a strong up-regulation of the Sperm-Leucylaminopeptidase 3 (S-lap3) compared to *Lp*^NIZO2877^-associated larvae, reaching a variation of 23-fold. It is important to notice that *Lp*^FLYG2.1.8^ triggered the up-regulation of five S-lap genes (*S-lap1*, *S-lap2*, *S-lap3*, *S-lap4*, *S-lap7*, *S-lap8*) compared to axenic larvae, but only *S-lap3* expression was significantly different compared to *Lp*^NIZO2877^-associated larvae (Table S1). Sperm-Leucylaminopeptidases have been described as structural components of the paracrystalline material of Drosophila sperm, and they collectively represent the most abundant protein class in Drosophila sperm (62). It has been shown that S-lap proteins are expressed exclusively during spermatogenesis (42). However, although gonadal development begins during larval and pupal stages, Drosophila larvae are not sexually mature (63–65). In fact, sexual maturation in Drosophila may require up to several weeks, depending upon the species. Here, we do not expect that the S-lap regulation is directly linked to fly fertility or sperm function. Instead, being peptidases, they may be involved in host development and maturation. This is in line with the up-regulation of protease genes that have been detected in presence of *Lp*^FLYG2.1.8^ (e.g., CG43124, CG43125). Beneficial microbes have been directly associated with increased proteolytic activities, which improve host nutrition and lead to faster development (46, 48). Here, we speculate that the up-regulation of Sperm-Leucylaminopeptidases exerted by *Lp*^FLYG2.1.8^ might impact larval nutrition and development, which is known to affect the subsequent adult reproductive success (66, 67). Indeed, *Lp*^FLYG2.1.8^ also triggers the up-regulation of genes directly involved in Drosophila development and lifespan (i.e., *mthl6*) and nutrient sensing (*Obp56e*). *mthl6* belongs to the Methuselah/Methuselah-like (Mth/Mthl) gene family of G protein-coupled receptors, which are essential for larval development, stress resistance, and in the setting of adult lifespan (68–70). On the other hand, the odorant-binding protein *Obp56e* is required for the detection of several essential amino-acids. Its up-regulation in the presence of *Lp*^FLYG2.1.8^ might be thus linked to a higher efficiency in nutrient sensing and assimilation as well as potentially higher food intake. Further work would be needed to investigate the actual role of S-lap family in larval maturation and growth and the consequences of altered expression on fly fertility.

The most surprising finding of this study was the down-regulation of the immune response exerted by the beneficial strain *Lp*^FLYG2.1.8^. Although, as mentioned above, both *Lp*^FLYG2.1.8^ and *Lp*^NIZO2877^ strains cause an increased expression of peptidoglycan recognition proteins compared to axenic larvae, the association with *Lp*^FLYG2.1.8^ resulted in the specific lower expression of *PGRP-Sc1a* and *PGRP-Sc1b* compared to *Lp*^NIZO2877^-associated larvae. The PGRP-SC gene cluster contains three PGRP-SC isoforms: PGRP-SC1a, PGRP-SC1b and PGRP-SC2. PGRP-SC1a and PGRP-SC1b probably arose from a recent duplication given that the two genes differ only by a synonymous mutation and produce the same polypeptide (28). They display an amidasic activity as they can bind and hydrolyse the bacterial peptidoglycan into smaller non-immunogenic muropeptides (45, 71–73). In contrast, non-catalytic PGRPs (i.e., PGRP-SA, PGRP-SD, PGRP-LC, PGRP-LE, etc.) bind to peptidoglycan but lack amidase activity. They are essential sentinels upstream of the two NF-κB-dependent *Drosophila* signalling cascades called Toll and immune deficiency (IMD) (58). While the mode of action of non-catalytic PGRPs in the Drosophila immune response and bacterial recognition has been extensively studied (74–84), the precise role of PGRP-SCs is less well defined. In this context, few studies have focused on determining the function of PGRP-SCs. Specifically, the amidasic activity of PGRP-SCs has been shown to result in dampening the IMD pathway, which appears to be under strong negative regulation in flies (28, 45, 85). This is known as immune tolerance, and it is meant to prevent constitutive activation of the energy-consuming NF-kB pathways (86). In addition, it has been demonstrated that PGRP-SC1 and PGRP-SC2 are required *in vivo* for full Toll pathway activation by Gram-positive bacteria and for phagocytosis (45, 87). Here, we reveal that a beneficial *L. plantarum* strain is able to exert its growth-promoting effect by specifically down-regulating PGRP-SC1, with no effect on other PGRPs. First, we hypothesised that such effect might be driven by the evolutionary history of *Lp*^FLYG2.1.8^, which has been co-evolving with the fly for two generations (39). However, our testing of other beneficial strains with different evolutionary backgrounds (i.e., *Lp*^DietG20.2.2^) led to the same effect in *PGRP-SC1* transcriptional regulation (Fig. 2A). In this regard, we demonstrated that the lower expression of *PGRP-SC1* directly results from the loss of function of the *L. plantarum ackA* gene. Indeed, we show that deleterious mutations in the *ackA* gene of different *L. plantarum* strains cause PGRP-SC1 down-regulation, which in turns leads to the improvement of the *L. plantarum* growth-promoting effect. It is interesting to note that the lower expression of *PGRP-SC1* does not result from decreased bacterial loads in the food or in the larval intestine. Contrarily, all *L. plantarum* strains bearing the *ackA* mutation exhibited higher loads compared to *Lp*^NIZO2877^. At the same time, we show that the bacterial metabolic rewiring following *ackA* mutation is sufficient to down-regulate the expression of PGRP-SC1. In particular, the supplementation of N-acetyl-glutamine, a direct consequence of *ackA* loss-of-function, was sufficient to trigger the lower expression of PGRP-SC1 without causing an increase in bacterial colony counts. This suggests that other metabolic pathways separate from those that increased NAG production are related to the improved growth of *L. plantarum* in association with Drosophila.

We next asked whether the down-regulation of PGRP-SCs was sufficient to improve the beneficial effect of *Lp*^NIZO2877^. Remarkably, we found that in *PGRP-SC*^*Δ*^ flies, *Lp*^NIZO2877^ significantly improves its benefit, recapitulating the effect exerted by *Lp*^FLYG2.1.8^. Importantly, we show that such improvement in growth promotion likely results from the higher bacterial loads reached by *Lp*^NIZO2877^ in *PGRP-SC*^*Δ*^ flies, compared to the reference population (Fig. 3B). Again, this result points out the importance of bacterial colony counts in affecting larval growth and development. Altogether, our data demonstrate that the degree of benefit exerted by *L. plantarum* towards Drosophila relies on the regulation of its immune response. Specifically, highly beneficial *L. plantarum* strains improve Drosophila growth by down-regulating the expression of PGRP-SC1 compared to moderate beneficial strains. The down-regulation of such peptidoglycan catalytic enzymes likely results in a decreased bactericidal activity and higher viable bacterial cells, which in turn benefit larval development and maturation. It is important to note that we did not detect any significant difference in the regulation of the IMD pathway (i.e., *pirk*, *Rel*, *AttD* gene expression) nor the Toll pathway (i.e., *Drs* expression) between *Lp*^FLYG2.1.8^- and *Lp*^NIZO2877^-associated larvae, indicating that the higher expression of *PGRP-SC1* observed in the presence of *Lp*^NIZO2877^ does not result in the IMD negative regulation, as expected from previous works (45, 85). Instead, it might directly lead to a higher bactericidal activity and the consequent decrease of bacterial loads. On the other hand, the increase in bacterial loads of *Lp*^FLYG2.1.8^ following PGRP-SC downregulation leads to improved host growth, further proving that bacterial concentration is directly proportional to microbial benefit. This is in line with the dual effect that commensal bacteria exert on their host. First, Drosophila larvae can use the microbial biomass as a source of additional nutrients, especially under nutrient-scarce conditions (88, 89). Secondly, live microbes can improve animal metabolism and amino acid absorption by increasing the host’s intestinal peptidases activity shown here and in prior work (51, 52). Moreover, live microbes actively produce and release essential nutrients (39, 88, 91, 92) whose density arises as bacterial loads increase. In this light, we specifically demonstrated that the metabolic products of live bacteria (i.e., N-acetyl-glutamine) are sufficient to trigger a lower expression of PGRP-SC1 and improve larval growth. The effect of bacterial metabolites on the regulation of Drosophila immune system has been also reported in previous studies (10, 93). Kamareddine et al. demonstrated that acetate produced by commensal bacteria from the *Acetobacter* genus can alter the expression of IMD pathway genes through PGRP-LC regulation in the enteroendocrine cells, which leads to mobilization of lipid resources in the nearby enterocytes and ultimately to growth promotion (93). While these data show that local signalling of bacterial by-products have systemic consequences on the host’s development, it remains unknown how such metabolites (i.e., N-acetyl-glutamine, acetate, etc.) directly affect Drosophila immune response.

Collectively, our work shows that the lower expression of PGRP-SC1 in Drosophila caused by beneficial commensal bacteria is directly responsible of improved larval growth. PGRPs are highly conserved from insects to mammals. Although, in insects, PGRPs are mostly involved in regulating defence pathways, in mammals, they have primarily antimicrobial activities (28, 94–96). Here, we show that a higher expression of PGRP-SC1 is directly linked to lower microbial loads, suggesting the specific bactericidal activity of such protein, with no effect on the regulation of immune pathways. This demonstrates that intestinal immune tolerance mechanisms to beneficial bacteria are critical to lead to higher bacterial growth and thus to improved host’s health. Signalling through immune pathways happens via the activity of bacterial peptidoglycan and metabolic products (i.e. N-acetyl-glutamine), which are directly responsible of host immune regulation and growth. Our results reinforce the notion that the influence of beneficial microbes on Drosophila physiology therefore relies on the intricate network of nutritional, metabolic and immune inputs (86). Further work would be needed to dissect the interdependency of such processes and to reveal the mechanisms underlying the bacterial specificity towards Drosophila immune effectors.

## MATERIALS AND METHODS

### Bacterial strains and culture conditions

The strains used in the present study are listed in Table S5. All strains were grown in Man, Rogosa, and Sharpe (MRS) broth and agar medium (Condalab, Spain) over night at 37°C without shaking and stored at −80°C in MRS broth containing 20% glycerol.

### *D. melanogaster* strains and maintenance

*Drosophila yw* flies were used as the reference strain in this work. The *PGRP-SC*^*Δ*^ KO line (kind gift from Prof. Bruno Lemaitre) is described in Paredes et al, (28). Germ-free (GF) stocks were established as described in Storelli et al. (34). *Drosophila* stocks were kept at 25°C with 12/12 hrs dark/light cycles on a yeast/cornmeal diet containing 50g/l inactivated yeast (rich diet) as described by Storelli et al. (34). Poor Yeast Diet (PYD) and PYD + N-acetyl-Glutamine (NAG) were obtained by reducing the amount of yeast extract to 8g/L and adding 1 g of NAG, respectively, as described in Martino et al. (39).

### Colonization of larvae and larval size measurements

GF adults were put overnight in breeding cages to lay eggs on PYD. GF embryos were collected the next morning and seeded by pools of 40 on 55mm petri dishes containing PYD. Bacterial strains were cultured to stationary phase in MRS broth at 37°C. The embryos were mono-associated with 150 μl (7×10^7^ total CFU) of the respective bacteria, or sterile Phosphate buffered saline (PBS, Sigma-Aldrich, USA) for GF condition. Petri dishes were incubated at 25°C until larvae collection. Images of the larvae were captured using a digital microimaging Leica DMD108 and larval longitudinal size (length) was measured as described in Martino et al (39).

### Bacterial load quantification

Quantification of larvae-associated bacteria was determined from a pool of 25 larvae (at least three replicates for condition) on day 4. Larvae were surface sterilized in 75% ethanol and placed in 1 ml of PBS. Samples were mechanically crushed, plated on MRS agar and incubated at 37°C for 48h. For each samples, the remaining fly food was placed in 10 ml of PBS. Samples were plated and incubated as reported above. The CFU count was performed using automatic colony counter Scan^®^ 300 (Vetrotecnica, Italy) and its accompanying software. GF samples were plated on Luria Bertani (LB) agar (Condalab, Spain), and incubated for 72h at 25°C.

### *L. plantarum*^NIZO2877^ mutant generation

#### Plasmid generation

Two *E. coli-Lactiplantibacillus* shuttle vectors were used to perform genome editing in *Lp*^NIZO2877^. The first shuttle vector, pCB578 (Table S6) encodes SpyCas9, a tracrRNA and repeat-spacer-repeat array with a 30-nt spacer targeting the acetate kinase gene (*ackA*) that was used previously in *Lp*^NIZO2877^ (39). The second shuttle vector, pCB591, was used to clone a recombineering template to generate a clean deletion of the *ackA* gene in *Lp*^NIZO2877^. First, pCB591 was amplified with primers oJV1-2, and genomic DNA from *Lp*^NIZO2877^ was amplified with primers oJV3-4 that amplify ackA as well as 1-kb homology arms flanking the start and stop codons. The PCR fragments were stitched together using the Gibson assembly kit (NEB CN# E2611S) following manufacturer’s instructions to yield pJV204. Then, primers oJV5-6 were used to remove *ackA* from pJV204 using the Q5 site-directed mutagenesis kit (NEB CN# E0554S) following the manufacturer’s instructions, yielding the final recombineering template pJV218. NEB 5-alpha or 10-beta Competent *E. coli* cells were used for both cloning steps, and primers oJV7-8 were used for screening clones by colony PCR. Correct clones were confirmed by Sanger sequencing.

#### Transformation of plasmids

Transformation of plasmids into *Lp*^NIZO2877^ was performed as described previously (98). In short, 1 ml of an overnight culture was used to inoculate 25 ml of fresh MRS supplemented with 2.5% glycine and was grown until OD600 reached 0.6-0.8. Then, cells were washed twice with ice-cold 10 mM MgCl_2_ and twice more with ice-cold SacGly solution. Plasmid DNA (1 μg suspended in water) and 60 μL of electrocompetent cells were added to a pre-cooled 1-mm electroporation cuvette and transformed at the following conditions: 1.8 kV, 200 Ω resistance, and 25 μF capacitance. Following electroporation, cells recovered in MRS broth for 3 hours then were plated on MRS agar containing appropriate antibiotics for 3 days before screening or inoculating colonies. Chloramphenicol and erythromycin concentrations were both 10 μg mL^−1^ in MRS liquid media and agar.

#### Genome editing

To delete *ackA* from *Lp*^NIZO2877^, the recombineering template shuttle vector pJV218 was passaged through the methyltransferase-deficient *E. coli* strain EC135 to improve transformation efficiency (99) and then transformed into *Lp*^NIZO2877^, where transformants were plated on MRS agar plates containing chloramphenicol. Then, the ackA-targeting SpyCas9 shuttle vector pCB578 was passaged through EC135 and transformed into the *Lp*^NIZO2877^strain harbouring the recombineering template shuttle vector, where transformants were plated on MRS agar containing erythromycin. Surviving colonies were screened for the desired genomic deletion using colony PCR with primers oJV9-10, and the PCR products were subjected to gel electrophoresis and Sanger sequencing with oJV11 to validate the clean deletion of *ackA* (Fig. S3). Both plasmids were cured from the mutant *L. plantarum^ΔackA^* strain by cycling between culturing in MRS media and plating on MRS agar, both without antibiotics. After each round of non-selective growth, cultures were plated on MRS agar supplemented with either chloramphenicol or erythromycin. This cycle was repeated until the mutant strain was sensitive to either antibiotic.

### RNA extraction, RT-PCR and Real-time PCR

Five biological replicates were generated for each condition (*Lp*^NIZO2877^-, *Lp*^Fly.G2.1.8^-, and PBS-associated larvae). Twenty-five larvae were collected for each sample, transferred in 50 μl of RNA later, flash-frozen in liquid nitrogen and stored at −80°C. RNA was extracted using RNeasy Mini Kit (Qiagen, Germany). Quantity and quality of RNA was assessed on an Agilent 2100 Bioanalyzer using Agilent RNA 6000 Nano kit (Agilent, USA). Extracted RNA was reverse-transcribed to cDNA using SuperScript™ IV First-Strand Synthesis System (Invitrogen™, USA). Real-time PCR amplifications were performed on a LightCycler 480 thermal cycler (Roche Diagnostic, Mannheim, Germany) in a final volume of 10 μl, which included 2.5 μl of cDNA template. The PowerUp™ SYBR™ Green Master Mix (Applied Biosystems™, USA) was used together with 0.25 μl of each primer (Table S7). The cycling conditions were as follows: 50 °C for 2 min, followed by 2 min at 95 °C, and 45 cycles at 95 °C for 10 s and 60°C for 1 min.

### Library preparation and RNA sequencing

The preparation of mRNA-Seq libraries and their sequencing has been carried out at the EMBL Genomics Core Facilities, Germany. Preparation of barcoded stranded mRNA-Seq libraries was performed with unidirectional deep sequencing of pooled libraries, read-length 80 bases, Illumina NextSeq run (yield ~550 million reads/lane), the pool of 15 libraries in one run.

### Bioinformatic and Statistical analyses

Single-end reads have been mapped onto the *D. melanogaster* reference dmel-all-r6.36 with STAR. The read counts were obtained with RSEM and the differential expression analysis has been done on R with DEseq2 package. For the comparative analysis between gnotobiotic and GF larvae, all samples and the three conditions were added to the model. To compare *Lp*^NIZO2877^- and *Lp*^Fly.G2.1.8^-associated larvae samples, a second model was used with only these samples. The GO term analysis was performed using DAVID online tool on the Flybase gene ID (41).

Data representation and analysis was performed using Graphpad PRISM 9 software (www.graphpad.com). One–Way ANOVA test was applied to performed statistical analyses between multiple (n>2) conditions for multiple comparisons. Additional images have been done on R software with ggplot2 package.

## Data Availability

Raw data (FASTQ) have been deposited at NCBI under the temporary Accession number SUB10001912.

## Author Contributions

M.G. and M.E.M designed the project; M.G. and E.M. conducted the experiments; P.J. and A.Q. conducted the bioinformatics analyses; J.V and C.B. designed and performed the CRISPR-Cas9 editing experiments; M.G. and M.E.M analysed the data and wrote the paper.

## Acknowledgements

Research in MEM lab was supported by the STARS@UNIPD Funding Programme (Starting Grant 2017 D.R. 872/2017) of the University of Padova. This work was also supported by the National Science Foundation (MCB-1452902 to CLB).

---

**FIG S1**. Comparison among the longitudinal size (mm) of *Lp*^NIZO2877^-, *Lp*^FlyG2.1.8^ and *Drosophila* germ-free (GF) larvae, respectively. Lines above each bar indicate the standard deviation (SD) calculated on N≥60 larvae analyzed for each condition.

**FIG S2** (A) Venn diagram – Number of genes with significantly enriched (up) or depleted (down) expression in gnotobiotic larvae (L) relative to axenic larvae (p < 0.05 and −1.5 to +1.5 fold). (B) GO terms (Biological Process) that were significantly represented (*Benjamini-adjusted-p-value* < 0.1) in the differentially expressed genes in gnotobiotic larvae relative to axenic larvae. (C) GO terms (Biological Process) that were significantly represented (*Benjamini-adjusted-p-value* < 0.1) in the differentially expressed genes in larvae associated with *Lp*^FlyG2.1.8^ relative to axenic larvae.

**FIG S3.** CRISPR-Cas9 genome editing in *L. plantarum*^NIZO2877^ to delete the *ackA* gene. (A) Scheme to generate a clean knockout of the *ackA* gene using the recombineering template and SpyCas9 shuttle vectors. First, shuttle vector pJV218 harboring a recombineering template containing 1000-bp homology arms (HA) on either side of the *ackA* coding region was transformed into *L. plantarum*^NIZO2877^. Then, the shuttle vector pCB578 encoding SpCas9, its tracrRNA, and a single-spacer CRISPR array targeting a site within *ackA* was transformed into the recombineering template strain to remove any unedited cells. Surviving colonies were screened by cPCR that amplifies from the genome of *L. plantarum*^NIZO2877^ but not from the plasmid with the recombineering template. (B) Validation of the *ackA* deletion via PCR and Sanger sequencing. Samples 1-2 on the gel show the PCR product obtained from the WT genome, while Samples 3-4 show the PCR product obtained from a mutant (A1) strain. The PCR product from the mutant strain was then subjected to Sanger sequencing with primer oJV11 to confirm that the start codon through the stop codon of *ackA* was successfully deleted.

## Tables – Legends

**Table S1**. RNA-seq data – Microbiota-dependent changes in average expression were analysed using DESeq package. The table shows Gene ID, mean, fold change expression, p-adjusted value and gene name in FlyBase format.

**Table S2**. Gene Ontology enrichment, full sets of terms and differentially expressed genes for (a) molecular function enriched in gnotobiotic larvae, (b) molecular function depleted in gnotobiotic larvae (c) molecular function depleted in *Lp*^FlyG2.1.8^-associated larvae, (d) molecular function enriched in *Lp*^FlyG2.1.8^-associated larvae, all relative to axenic larvae.

**Table S3.** List of genes differentially expressed (up- and down- regulated) between Lp^FlyG2.1.8^- and Lp^NIZO2877^-associated larvae, selected for values of p < 0.05 and −1.5 to +1.5 fold. The table shows Gene ID, gene name in FlyBase format, mean, fold change expression and p-adjusted value.

**Table S4**. List of molecular function derived from UniProt database associated with the genes differentially expressed (up- and down-regulated) between Lp^FlyG2.1.8^- and Lp^NIZO2877^-associated larvae. (https://www.uniprot.org)

**Table S5**. Bacterial strains used in the present work

**Table S6**. Plasmids used in the present work.

**Table S7**. Sequences of primers used in the present work.

