## Supplementary figures and images for "Beneficial *Lactiplantibacillus plantarum* promote Drosophila growth by down-regulating the expression of PGRP-SC1"

### Supplemental Figure 1

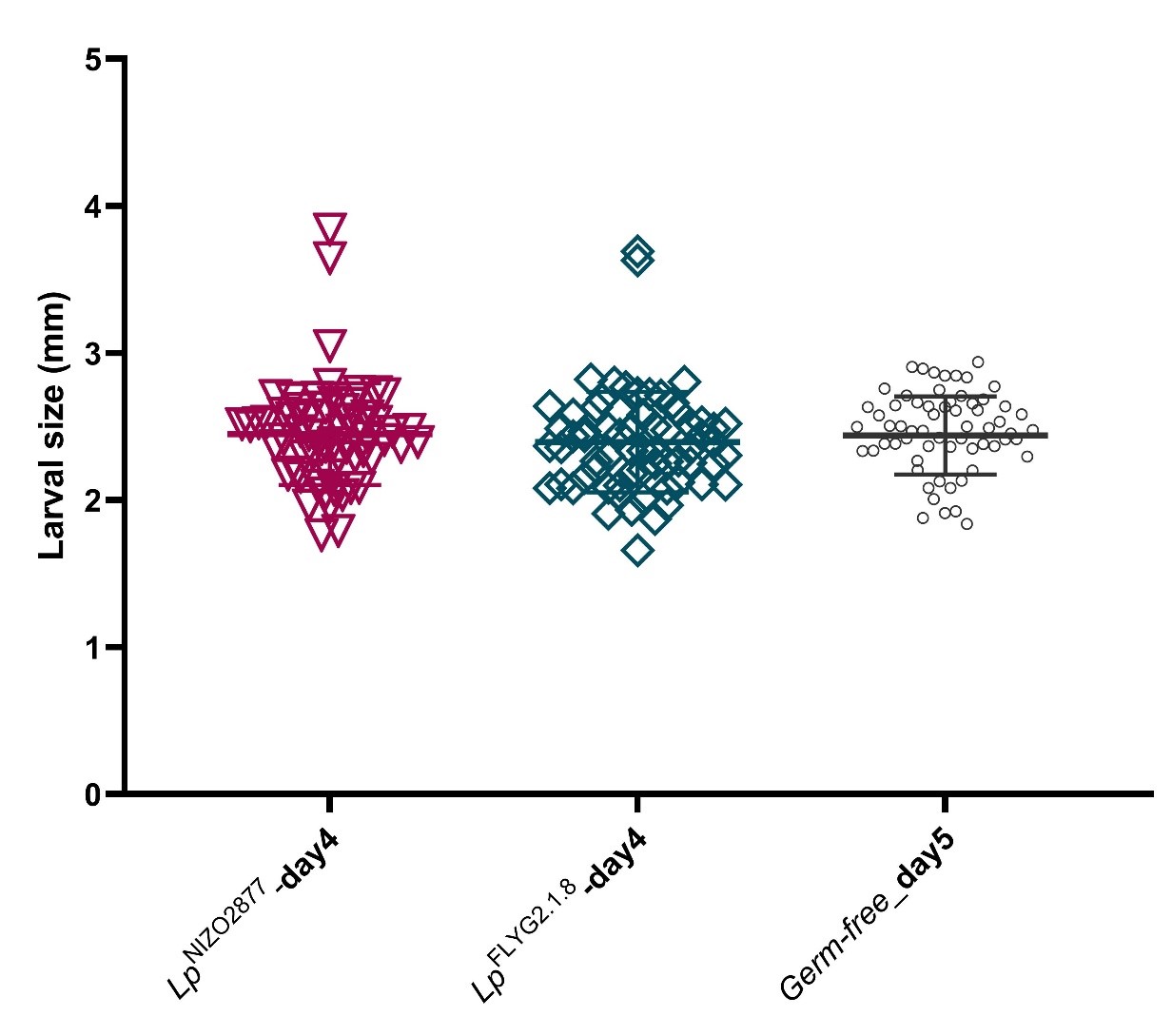

### Supplemental Figure 2

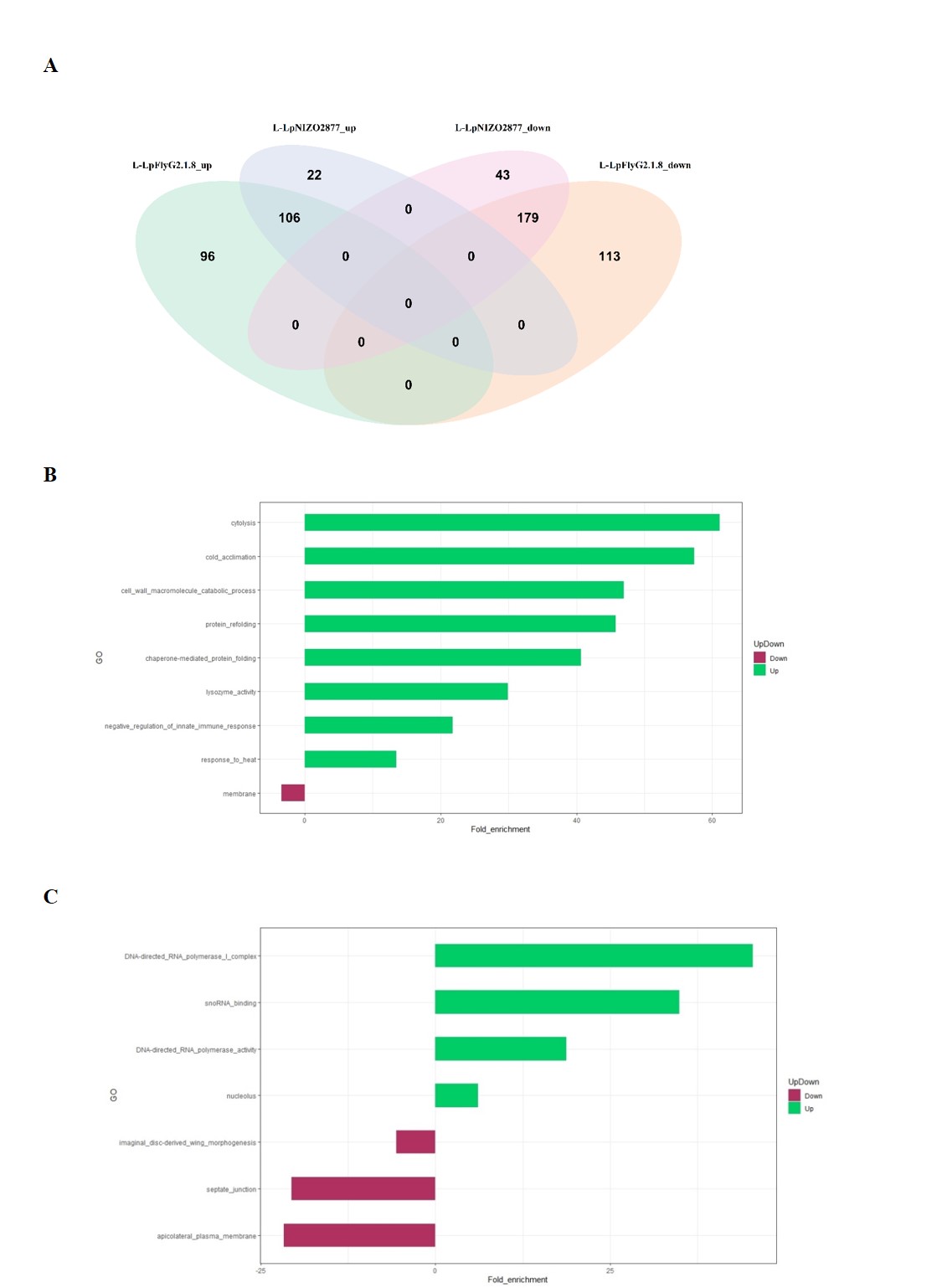

### Supplemental Figure 3

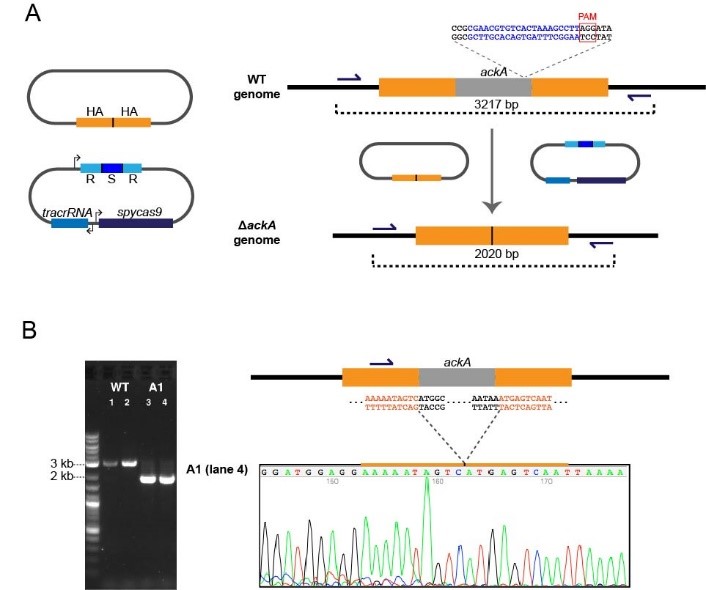
